# Pocket-Conditioned Generation and Down-Selection of Non-Oxime Reactivators for Paraoxon-Inhibited Human Acetylcholinesterase

**DOI:** 10.64898/2026.10.09.758041

**Authors:** Daniel Jimenez, William K. Clay, Carlos Barajas

## Abstract

Acetylcholinesterase reactivation is a structure-constrained design problem in which chemical plausibility, access to central nervous system, pocket compatibility, and reactivation geometry must be considered together. We report a computational workflow for generating and down-selecting candidate non-oxime reactivators in the paraoxon-inhibited human acetylcholinesterase pocket. For generating the candidates, the first-round utilitezed pocket-conditioned DrugFlow, but due to solubility issues, the second-round candidates were output form FlexFlow, DrugFlow-OOD, DrugFlow preference alignment, and pocket-specified BoltzMol-1. A property-guided DrugDiff output provided a protein-unconditioned baseline. Although 93.9% of the DrugDiff subset passed Lipinski criteria for druglikeness, its median synthetic accessibility (SA) score signalled that it was difficult to synthesize compared against other generative approaches. A strict predicted blood-brain barrier and solubility gate retained 31.0% of Base DrugFlow, 61.0% of FlexFlow, 27.5% of DrugFlow-OOD, 3.8% of DrugFlow-PA, and 35.1% of BoltzMol-1. The second-round top-ten lists had lower SA scores and higher quantitative estimate of drug-likeness (QED) values than the DrugDiff shortlist, while DrugDiff had fewer aggregate heuristic violations. This work compares the best performing generative drug discovery models with suggested down sleection approaches for human acetylcholinesterase reactivation.

## 1 Introduction

Organophosphorus compounds inhibit human acetyl-cholinesterase (AChE) by covalently phosphorylating the serine residue of the catalytic triad. Inhibition of AChE leads to a buildup of acetylcholine which causes a cholinergic crisis and death [1, 2]. Current oxime antidotes can restore AChE activity for some inhibited states, but performance depends on multiple factors, including the organophosphorus compound structure, OP-inhibited AChE aging rate, and OP-inhibited AChE pocket geometry [3–7]. Quaternary oximes also have limited blood-brain barrier (BBB) penetration,[8, 9] a necessary characteristic to treat AChE in the brain and central nervous system. These constraints motivate alternative reactivator chemical classes [10] such as bis-pyridinium oxime scaffolds and other non-oxime scaffolds such as quinone methide precursors (Mannich bases) and imidazole-based compounds. Research on non-oxime reactivators has been recently reviewed [5, 11], and efforts on new reactivators continues [12, 13]. Overall, there is much unexplored chemical space for the development of non-oxime reactivators.

The structural use case, shown in Figure 1a, is Protein Data Bank entry 5HF9, a 2.20 Å structure of paraoxon-inhibited human acetylcholinesterase in complex with the oxime HI-6 [14, 15]. Paraoxon inhibition phosphorylates the active site serine residue, perturbs the acyl loop and narrows the gorge near the peripheral site. A small molecule generator that targets only bulk physicochemical properties cannot directly account for these geometric constraints. The design problem therefore requires protein or pocket conditioning, followed by property filtering and structural review.

**Figure 1:**
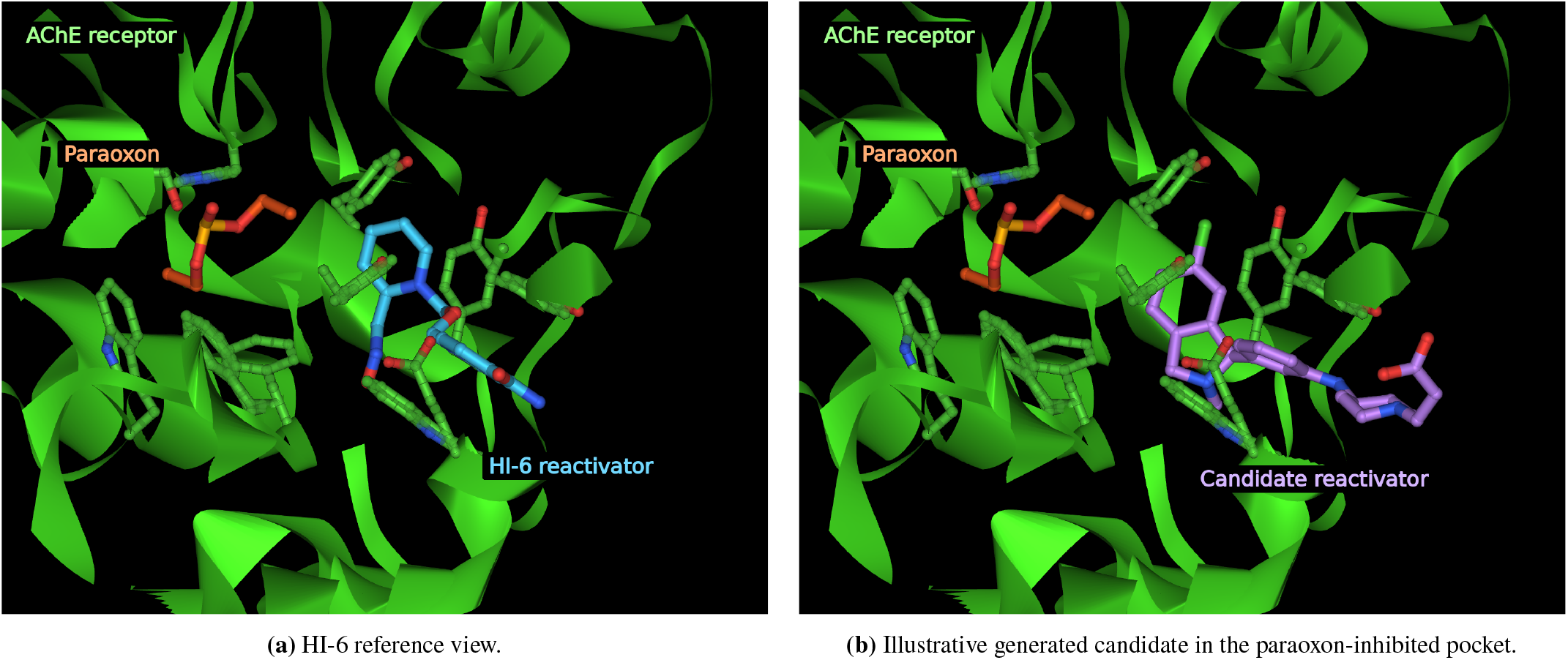
Paraoxon-inhibited human acetylcholinesterase structures used for pocket-conditioned candidate generation. Protein atoms and secondary structure are shown around the inhibited active site.

Property-guided generation remains useful as a comparison. DrugDiff uses a latent diffusion model with predictor guidance to steer molecular properties without retraining the diffusion model for each new objective [16]. Its generation process does not use the three-dimensional AChE pocket as context. DrugFlow generates atom types, bond types, and co-ordinates conditioned on protein structure [17]. Its extensions add binding-site side-chain sampling, an out-of-distribution uncertainty estimate, or preference alignment toward selected molecular objectives. The central question is whether structural conditioning, combined with explicit multi-stage selection, yields candidates that better satisfy the design specification than protein-unconditioned property guidance alone.

The analysis is limited to generated structures with their computed triage values based on the two-dimensional SMILES representation. These down-selection filters prioritize molecules for follow-up and do not establish AChE binding or biochemical reactivation.

## 2 Design Objective

The desired molecule is a non-oxime candidate that can access the paraoxon-inhibited AChE gorge, maintain a plausible conformation near the inhibited catalytic center, and satisfy early drug-like property constraints. These requirements are not reducible to a single score. Increasing quantitative estimate of drug-likeness (QED) or decreasing synthetic accessibility (SA) does not guarantee pocket complementarity, while optimizing a structural score can favor large, lipophilic, or poorly soluble molecules.

Figure 1 shows the known oxime, HI-6, and an AI-generated non-oxime candidate in the paraoxon-inhibited AChE pocket.

## 3 Computational Workflow

### 3.1 Target preparation and conditioning context

The campaign used the paraoxon-inhibited 5HF9 structure as the common structural reference. Protein coordinates and the inhibited-state context were prepared for target-aware generation, and the relevant pocket region was specified for each model interface. This shared context allowed for the comparison of multiple generation and screening strategies while keeping the biological objective fixed.

### 3.2 Generative and screening branches

We used two computational rounds. Each of the five pocket-aware branches targeted a pragmatic approximately 10^3^-molecule screen: large enough to quantify filtering attrition and extract a ten-candidate shortlist, but small enough to run across several structural variants.

#### Round 1: Base DrugFlow

DrugFlow samples ligand coordinates, atom types, and bond types while conditioning on fixed protein backbone coordinates and residue identity. Virtual nodes allow the final ligand size to vary. The first round used a permissive predicted-solubility gate that accepted moderately soluble candidates.

#### Round 2: Diversified target-aware branches

Four additional branches broadened the search while using a stricter solubility gate.

- **FlexFlow** jointly samples ligand variables and binding-site side-chain torsion angles χ. The protein backbone remains fixed, while pocket rotamers can change [17].
- **DrugFlow-OOD** adds an uncertainty head trained with the flow-matching objective. Per-atom uncertainty is integrated over the sampling trajectory to support confidence and out-of-distribution (OOD) assessment [17].
- **DrugFlow-PA combined** starts from the base DrugFlow weights and applies pairwise preference alignment (PA). The combined source objective favors higher QED, lower synthetic accessibility (SA), improved Vina efficiency, and REOS passage [17].
- **BoltzMol-1** uses the web interface https://lab.boltz.bio/ to create both a new target from the protein sequence and specified pocket residues as well as a new experimental screening configuration. The constraints restricted proposals to Enamine REAL Space, described in the run configuration as vetted and readily synthesizable compounds, and required Lipinski Rule-of-Five compliance and molecular weight below 500. The run requested virtual generation and screening of 1,000 candidates at a recorded total cost of $65.95. The downloaded export was restricted to molecules assigned *None* or *Minor* structural-alert classification by the platform, leaving 202 molecules for the common analysis. BoltzMol-1 applies curated SMARTS-based structural-alert filters to identify predefined molecular substructures associated with potential toxicity, chemical reactivity, or poor pharmacokinetic properties. The platform reports the exported structural-alert classifications as *None, Minor, Major, Severe*, or *Critical*.

#### Protein-unconditioned comparison: DrugDiff

DrugDiff was guided toward molecular properties but was not conditioned on the AChE pocket [16]. To keep this comparison on the same scale as the pocket-aware branches, the descriptor analysis used a fixed random subset of 1,000 unique molecules from the DrugDiff export. A separately ranked DrugDiff top-ten export was retained for shortlist comparisons.

### 3.3 Descriptor calculation and medicinal chemistry filters

Candidate SMILES were evaluated with RDKit-based code [18]. The common absorption, distribution, metabolism, and excretion (ADME) characterization tables report synthetic accessibility (SA), Delaney estimated log solubility (ESOL), qualitative solubility class, predicted blood-brain barrier permeability (BBB), Lipinski violations, PAINS and Brenk alerts, several rule-based filters, REOS, and QED [19–25].

The ESOL classes were assigned as follows: *Insoluble* for log *S* < −10, *Poorly* for −10 ≤ log *S* < −6, *Moderately* for−6 ≤ log *S* < −4, *Soluble* for −4 ≤ log *S* < −2, *Very* for−2 ≤ log *S* < 0, and *Highly* for log *S* ≥ 0 where log *S* is the estimated log solubility. These are model-derived classes rather than measured solubilities.

The field bbb_permeant is a binary BOILED-Egg prediction based on lipophilicity and topological polar surface area [26]. A value of Yes denotes predicted passive blood-brain barrier permeation. It is used as a computational eligibility gate, not as an experimental permeability measurement.

### 3.4 Deduplication, eligibility, and ranking

All SMILES were parsed and canonicalized with RDKit. Duplicate canonical SMILES were removed before any denominator, distribution, or selection yield was calculated, with the first occurrence retained. The selection procedure then separated hard eligibility gates from ranking criteria.

1. **Predicted BBB gate:** bbb_permeant=Yes.
2. **Round-1 solubility gate:** *Moderately, Soluble, Very*, or *Highly*. This is denoted *Moderate+*.
3. **Round-2 solubility gate:** *Soluble, Very*, or *Highly*. This is denoted *Soluble+*.
4. **Ranking:** ascending aggregate filter-violation count, ascending SA, and descending QED.

The aggregate filter-violation count was ranked first because the counted terms correspond to explicit medicinal-chemistry failure modes, whereas SA and QED provide continuous criteria for synthetic tractability and drug-likeness. SA was ranked before QED because experimental follow-up requires accessible compounds, while QED is an aggregate drug-likeness heuristic that partly overlaps the preceding filters. The stricter second round was introduced because all ten first-round selections were classified as only *Moderately* soluble by ESOL. Listing 1 reproduces the common logic.

#### Listing 1

Canonical-SMILES deduplication and selection logic.

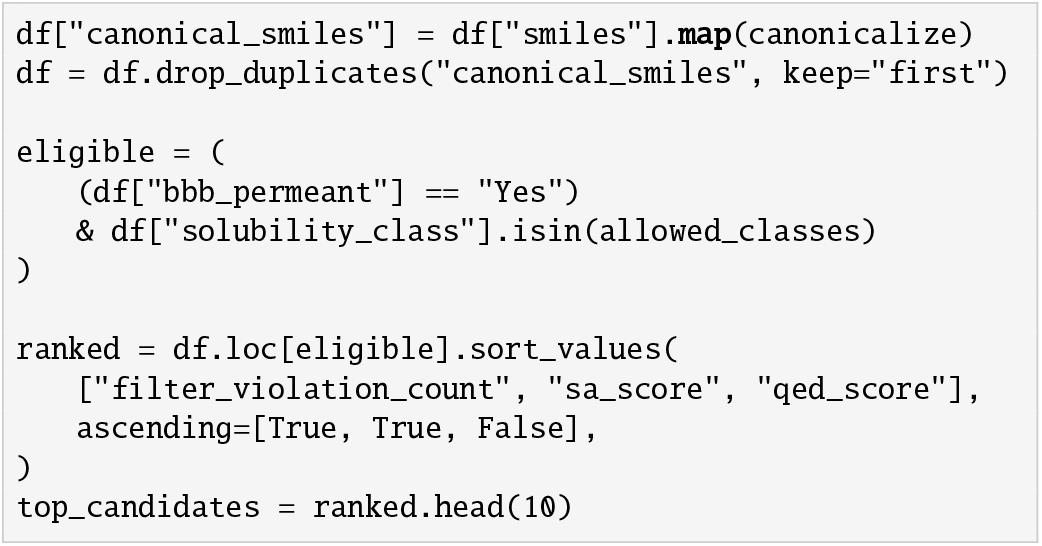

### 3.5 Interpretation of the aggregate violation count

The aggregate count is a reproducible heuristic profile, not a validated objective function. The implementation adds a Lipinski-failure indicator and a Rule-of-Five failure separately even though they use the same thresholds. It also counts failure of both the Rule of Four and Rule of Three. These rules describe distinct chemical regimes. The Rule of Four characterizes larger and more lipophilic protein-protein interaction inhibitors, while the Rule of Three describes fragment-like compounds [27, 28]. Their molecular-weight conditions do not overlap in the implementation, so a molecule cannot satisfy both. The Rule-of-Four and Rule-of-Three pass counts are therefore treated only as audit fields rather than as shortlist decision metrics.

The Ghose and Veber filters were included as early oral-drug heuristics [29, 30]. They guide review but do not replace target-specific medicinal chemistry assessment.

### 3.6 Diversity and non-oxime checks

Shortlist diversity was summarized with canonical SMILES, Bemis-Murcko scaffolds, and Morgan fingerprint Tanimoto similarity [31, 32]. Bemis-Murcko scaffolds are coarse chemical class representations that retain molecular ring systems and linkers while removing terminal side chains; counting them tests whether a shortlist spans multiple core architectures rather than repeated side-chain substitutions around one core. A preliminary oxime SMARTS screen of neutral oxime motifs, [CX3]=[NX2][OX2H1], demonstrated that none of the candidates matched this pattern. This pattern does not replace tautomer-aware enumeration or expert substructure review.

## 4 Results

### 4.1 Pool scale, deduplication, and descriptor distributions

The six analysis pools contained 4,554 candidates after canonicalization and deduplication. Every SMILES parsed successfully. DrugDiff was represented by 1,000 unique structures sampled from its canonicalized export. The five common ADME tables contributed 3,554 rows and 3,552 unique structures.

Table 1 compares properties present across all six analysis pools. DrugDiff had a 93.9% Lipinski pass rate, comparable with Base DrugFlow, FlexFlow, and DrugFlow-OOD. Its median SA was nevertheless 4.63, while the other pools ranged from 2.06 for DrugFlow-PA to 3.32 for FlexFlow. DrugDiff also had the lowest median QED at 0.441. The comparison shows that a high Lipinski pass fraction does not imply favorable synthetic accessibility or a strong QED distribution.

**Table 1:**
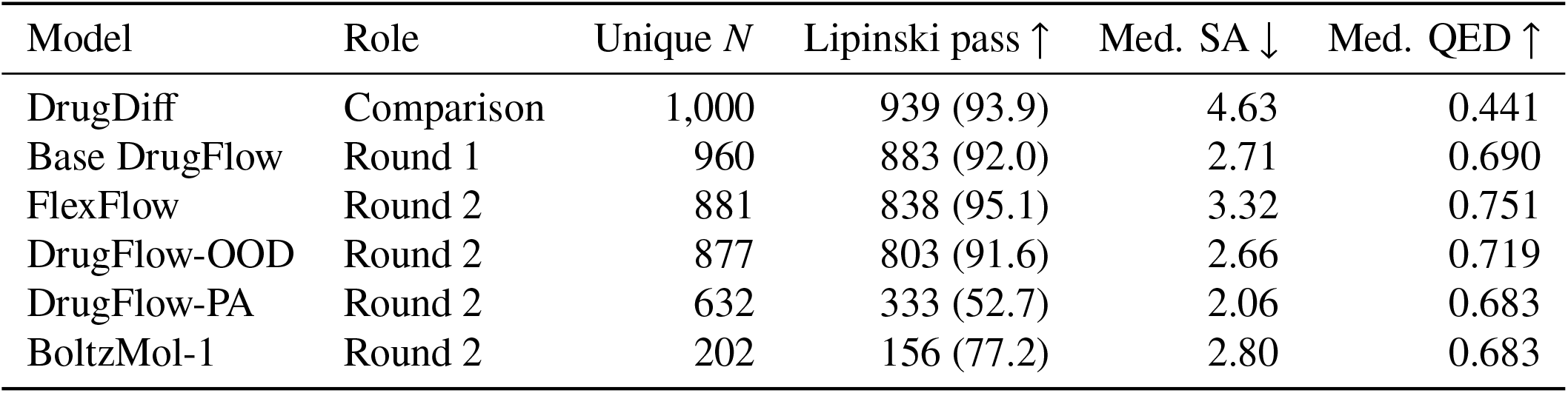
Analyzed-pool summary after canonical-SMILES deduplication. DrugDiff was represented by a fixed 1,000-molecule canonical-unique subset. Lipinski pass counts are followed by percentages. DrugDiff status was reconstructed from molecular weight, logP, hydrogen-bond donor, and hydrogen-bond acceptor columns; the other pools use their reported violation counts. SA and QED are medians over each unique pool. Arrows mark preferred directions. For example, molecules with more favorable drug-like properties are denoted with a higher QED score, where the QED score can range from 0 to 1.

### 4.2 Effect of the stricter solubility gate

Figure 2 reports eligibility under both solubility gates for the five outputs with common BBB and ESOL fields. Strict eligibility was highest for FlexFlow: 61.0% of its candidates passed both the predicted BBB gate and the *Soluble+* ESOL gate, compared with 35.1% for BoltzMol-1, 31.0% for Base DrugFlow, 27.5% for DrugFlow-OOD, and 3.8% for DrugFlow-PA. The preference-aligned strict subset had the lowest median SA and highest median QED among *Soluble+* candidates, but only 24 molecules satisfied both hard gates.

**Figure 2:**
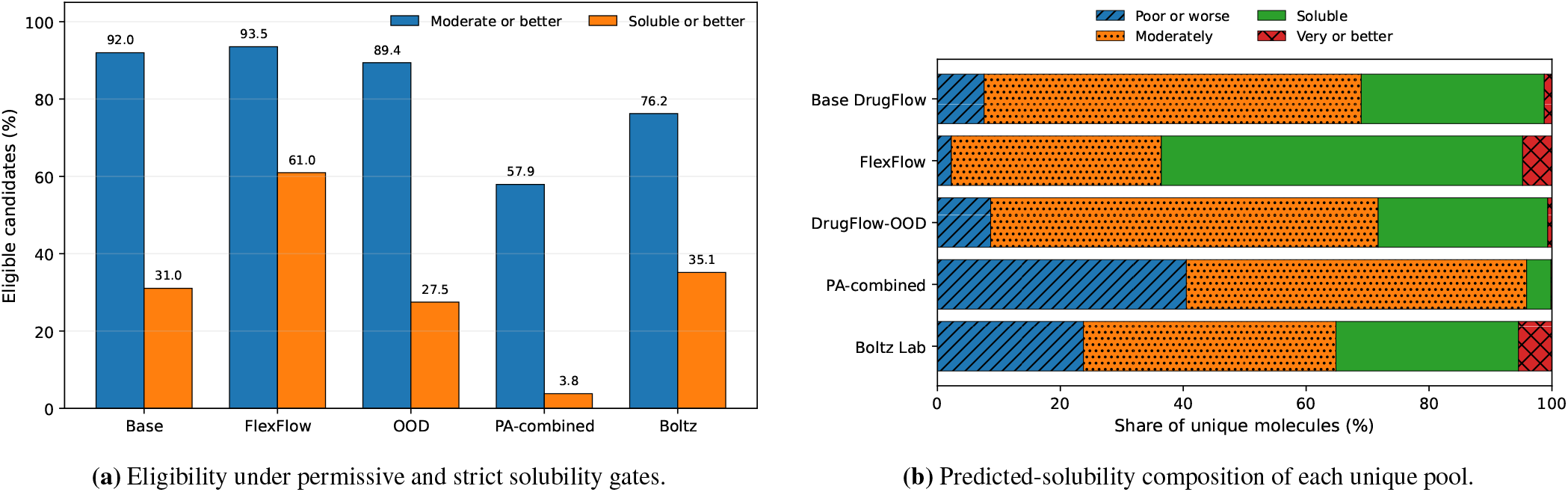
The stricter second-round gate changes model yield substantially. (A) Tightening the ESOL gate from *Moderate+* to *Soluble+* reduced yeild for every generative model, with FlexFlow retaining the largest fraction, BoltzMol-1 intermediate, Base DrugFlow and DrugFlow-OOD lower, and DrugFlow-PA lowest. (B) The same trend follows from predicted-solubility composition: FlexFlow and BoltzMol-1 contained the largest *Soluble*-or-better fractions, Base DrugFlow and DrugFlow-OOD contained larger *Moderately* soluble fractions, and DrugFlow-PA was dominated by *Moderately* and *Poorly* soluble molecules. Percentages use unique molecules as denominators.

The yield difference between these models is explained by the predicted-solubility composition in Figure 2. FlexFlow placed 58.8% of its unique molecules in the *Soluble* class and 4.8% in *Very* or *Highly* classes. The BoltzMol-1 export contained 29.7% *Soluble* and 5.4% *Very* molecules. DrugFlow-PA contained 55.4% *Moderately* and 40.5% *Poorly* soluble molecules, leaving 4.1% in *Soluble* or better before the BBB gate.

### 4.3 Shortlist tradeoffs

Table 2 compares six top-ten lists after their branch-specific hard gates. The Base DrugFlow list used the first-round *Moderate+* gate, while the DrugDiff and second-round target-aware lists used the stricter *Soluble+* gate. DrugDiff had a median aggregate violation count of one, but its median SA was 5.50, suggesting that the most drug-like molecules generated by DrugDiff would be difficult to synthesize. The four strict second-round target-aware lists had median SA values from 1.86 to 2.52 and median QED values from 0.796 to 0.865. DrugFlow-OOD had the lowest median SA, while FlexFlow had the highest median QED. BoltzMol-1 produced an intermediate profile with median SA 2.52 and median QED 0.796. These results show that a lower aggregate violation count did not imply better SA or QED.

**Table 2:**
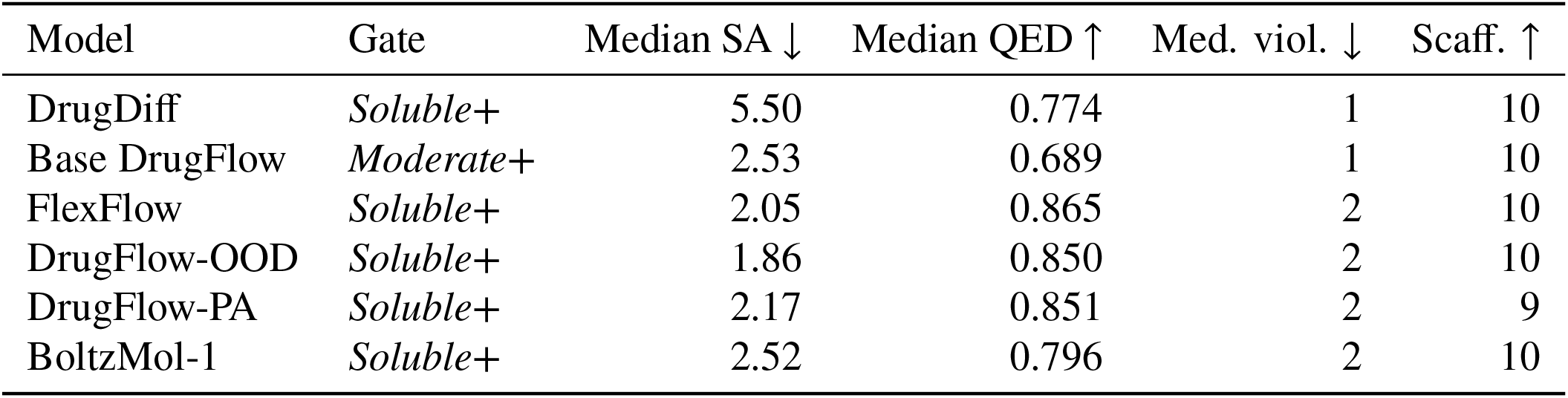
Summary of the six top-ten lists. Each row summarizes the metrics for the ten unique molecules selected with the gate shown. SA and QED are medians. Scaff. is the number of unique Bemis-Murcko scaffolds. Arrows mark preferred directions.

The Base DrugFlow shortlist had a median SA of 2.53 and median violation count of one, but all ten candidates were only *Moderately* soluble. That result motivated the stricter second round. Figure 3 visualizes the SA–QED tradeoff for the same 60 candidates. The second-round target-aware candidates occupy a lower-SA, higher-QED region relative to DrugDiff. This selected-set comparison does not establish binding or rank the full generators on a single universal objective. The ranked molecular structures for all six shortlists are provided in Section B.

**Figure 3:**
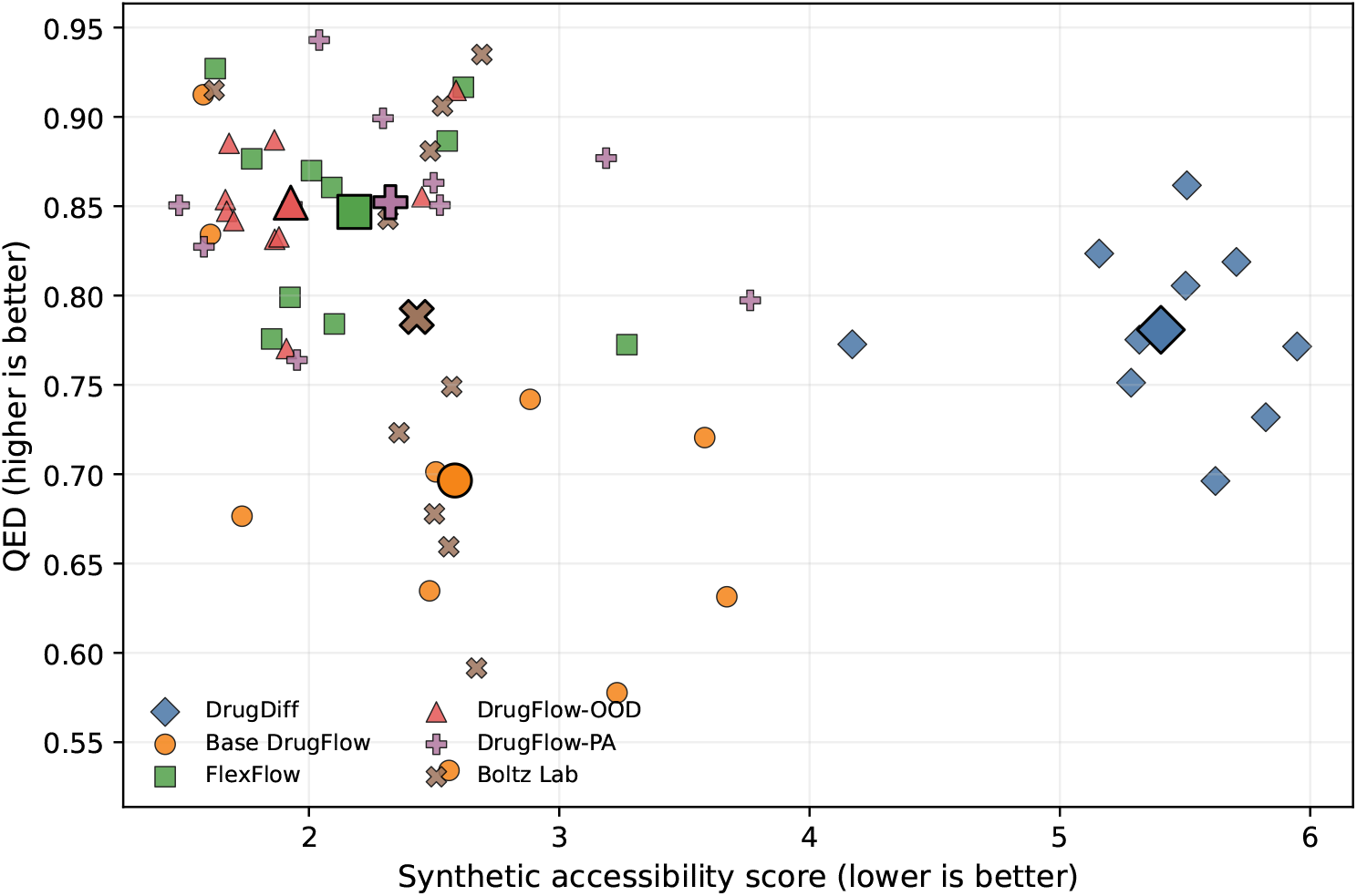
Synthetic accessibility and QED for the 60 shortlist molecules. Larger markers denote model means. Lower SA and higher QED are preferred.

### 4.4 Rule profiles, diversity, and non-oxime intent

All ten DrugDiff shortlist molecules failed the implemented Rule of Four and passed the Rule of Three. Seven of ten Base DrugFlow candidates passed the Rule of Four, while none passed the Rule of Three. The strict second-round lists all failed the Rule of Four. Rule-of-Three passage was 7 of 10 for FlexFlow, 2 of 10 for DrugFlow-OOD, 3 of 10 for DrugFlow-PA, and 2 of 10 for BoltzMol-1. These values describe molecular-size and lipophilicity regimes. Failing either rule was acceptable because the campaign did not screen fragments or explicitly seek large, lipophilic ligands for protein interfaces; the two rules were retained only to diagnose selected molecular size and lipophilicity.

All 60 shortlist molecules had unique canonical SMILES and represented 58 unique Bemis-Murcko scaffolds. Mean within-list pairwise Tanimoto similarity was between 0.12 and 0.19, indicating broad structural diversity at the shortlist scale. The preliminary oxime SMARTS screen found no matches.

## 5 Discussion

### 5.1 Conditioning is necessary but not sufficient

DrugDiff showed that property guidance can produce Lipinski-compliant molecules, but it sampled without the paraoxon-inhibited AChE gorge. The DrugFlow-family branches and BoltzMol-1 used pocket information, then the BBB and solubility gates removed different fractions from each pool. Base DrugFlow showed that pocket conditioning alone did not solve the filtering problem: its top ten had favorable SA values but only moderate predicted solubility.

Each second-round branch covered a different failure mode. FlexFlow relaxed the rigid-side-chain assumption and produced the largest strict eligible fraction. DrugFlow-OOD produced the lowest-SA shortlist. Preference alignment produced high-QED, low-SA candidates within a small strict subset. BoltzMol-1 added a constrained commercial-space screening route and retained 71 strict candidates from 202 exported molecules. We therefore retained candidates from all four branches rather than selecting a single generator.

### 5.2 Selection metrics should remain decomposable

The aggregate filter count is useful as a deterministic first sort key, but its current construction obscures interpretation. It double counts Lipinski-like failure and treats fragment-like and Rule-of-Four-like regimes as simultaneous requirements. A revised score should remove redundant terms or report a vector of component outcomes. Pareto ranking across SA, QED, predicted solubility, molecular size, structural alerts, model confidence, and interaction geometry would avoid collapsing partially conflicting criteria into the current violation count.

The DrugDiff subset illustrates this point. Its Lipinski pass fraction was high, but the SA and QED distributions were less favorable than those of the target-aware pools. Lipinski criteria are therefore useful exclusion rules, not a complete measure of design quality. Predicted BBB permeability and ESOL class should likewise be treated as computational hypotheses until measured.

### 5.3 Relationship to virtual screening and gated design workflows

Ligand-based tools such as VSFlow can rapidly search substructures, fingerprints, and three-dimensional shape once reference ligands or libraries are available [33]. Such tools can identify purchasable analogs of selected scaffolds and expand around high-priority chemical classes. They complement rather than replace target-conditioned generation.

Our workflow similarly generated broad candidate pools, applied common eligibility criteria, and retained structurally diverse shortlists for follow-up [34]. For small-molecule re-activation, the scorecard must keep chemistry-specific failure modes visible and connect computational prioritization to a functional assay, such as an Ellman reactivation assay [35].

## 6 Future Work and Compute Scaling

Future work might incorporate additional docking and structural analysis to prioritize candidates with favorable positions and orientations within the paraoxon-inhibited AChE pocket. Particular attention will be given to identifying nucleophilic or basic functional groups positioned to support either direct or water-mediated reactivation mechanisms. In the direct mechanism, the molecule would react with the inhibitory group attached to AChE, whereas in the water-mediated mechanism, it would facilitate the removal of that group through a reaction involving water. These structural assessments will help further narrow the candidate pool before experimental testing.

Further structural evaluation may use conventional all-atom protein-ligand molecular dynamics with established biomolecular force fields. Explicit-solvent simulations can test pose retention, pocket-side-chain rearrangements, and water-mediated interactions under controlled protonation and ionic conditions. Docking and molecular dynamics should be treated as complementary structural checks rather than substitutes for bio-chemical reactivation measurements. We evaluated PLACER [36] pRMSD because conformational ensemble predictions could provide a structure-aware confidence signal beyond the two-dimensional ADME and medicinal-chemistry filters. However, this exploratory analysis did not sufficiently distinguish the generated candidates to support primary ranking and is described in Section A.

Following computational down-selection, selected molecules will require synthesis and biochemical evaluation of AChE reactivation using Ellman’s assay. The experimental results will be reported in subsequent work.

## 7 Conclusion

This work demonstrates a reproducible workflow that combines pocket-conditioned small-molecule generation with physicochemical, medicinal-chemistry, and drug-likeness filters to prioritize diverse candidate non-oxime reactivators for paraoxon-inhibited human AChEN? on-oxime reactivators are of particular interest because existing oxime therapies can have limitations in efficacy and central nervous system exposure, leaving substantial chemical space to explore. However, generative models can produce large numbers of plausible molecules, making systematic down-selection necessary to reduce the experimental hypothesis space and focus time and resources on the most promising candidates. Rather than relying on a single generative model or scoring criterion, we evaluated multiple generation and down-selection strategies to identify candidates that satisfied the properties of interest. The resulting shortlists should be viewed as experimentally prioritized hypotheses rather than validated reactivators. Neither AChE binding nor functional enzyme reactivation has been established, and both will require experimental validation. The same general workflow could be adapted to other structure-conditioned small-molecule discovery problems by changing the target, generative model, and selection criteria. Separating generation from down-selection also makes it possible to incorporate assay-specific constraints as needed. We envision this approach being used to compare generative strategies and to narrow large generated chemical spaces to smaller sets of candidates suitable for tractable experimental follow-up.

## 8 Data Availability

The data underlying this study are available from the corresponding author upon reasonable request.

## 9 Acknowledgments

The authors acknowledge funding and computational resources from The Johns Hopkins University Applied Physics Laboratory Independent Research and Development Program. We also thank Dr. Laura Dunphy for helpful comments and suggestions that improved the clarity of the manuscript.

Boltz Bio platform credits were provided for beta testing. Large language model (LLM) tools were used to assist with software development, including code generation, debugging, refactoring, and documentation, as well as with drafting, editing, and improving the clarity of portions of the manuscript. All LLM-assisted code and text were reviewed, modified as needed, and validated by the authors, who take full responsibility for the final software, analyses, and manuscript content.

## Appendix

### A Exploratory PLACER Based Conformational Assessment

As a separate structural check, we ran PLACER to test whether the predicted RMSD (pRMSD) values from repeated predictions separated consistently modeled protein-ligand configurations from uncertain ones.

The generated candidates had uniformly low and tightly clustered pRMSD distributions. The HI6 reference control instead had a broader pRMSD distribution with a higher mean. Low pRMSD therefore did not separate the generated candidates or track the reference compound as expected, so we left PLACER pRMSD out of the primary ranking.

### B Top Candidate Molecular Structures

The following figures show the top 10 molecular candidates from each candidate set, in the same rank order as the corresponding top-picks CSV files. DrugFlow and DrugDiff were evaluated in Round 1; BoltzMol-1, FlexFlow, DrugFlow-OOD, and DrugFlow-PA were evaluated in Round 2.

**Figure 4:**
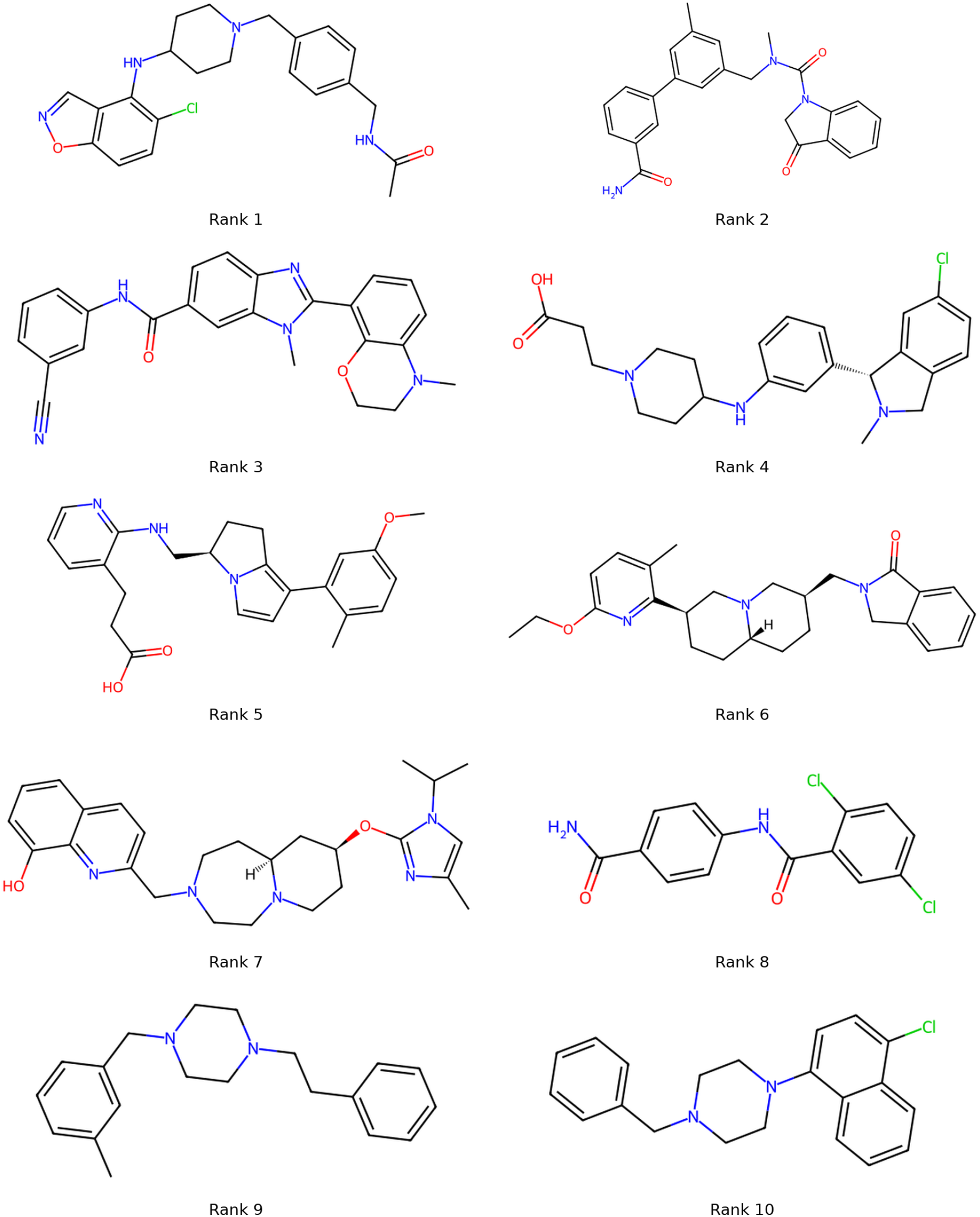
Top 10 DrugFlow candidate molecular structures from Round 1. Candidates are shown in the same rank order as the corresponding top-picks CSV file.

**Figure 5:**
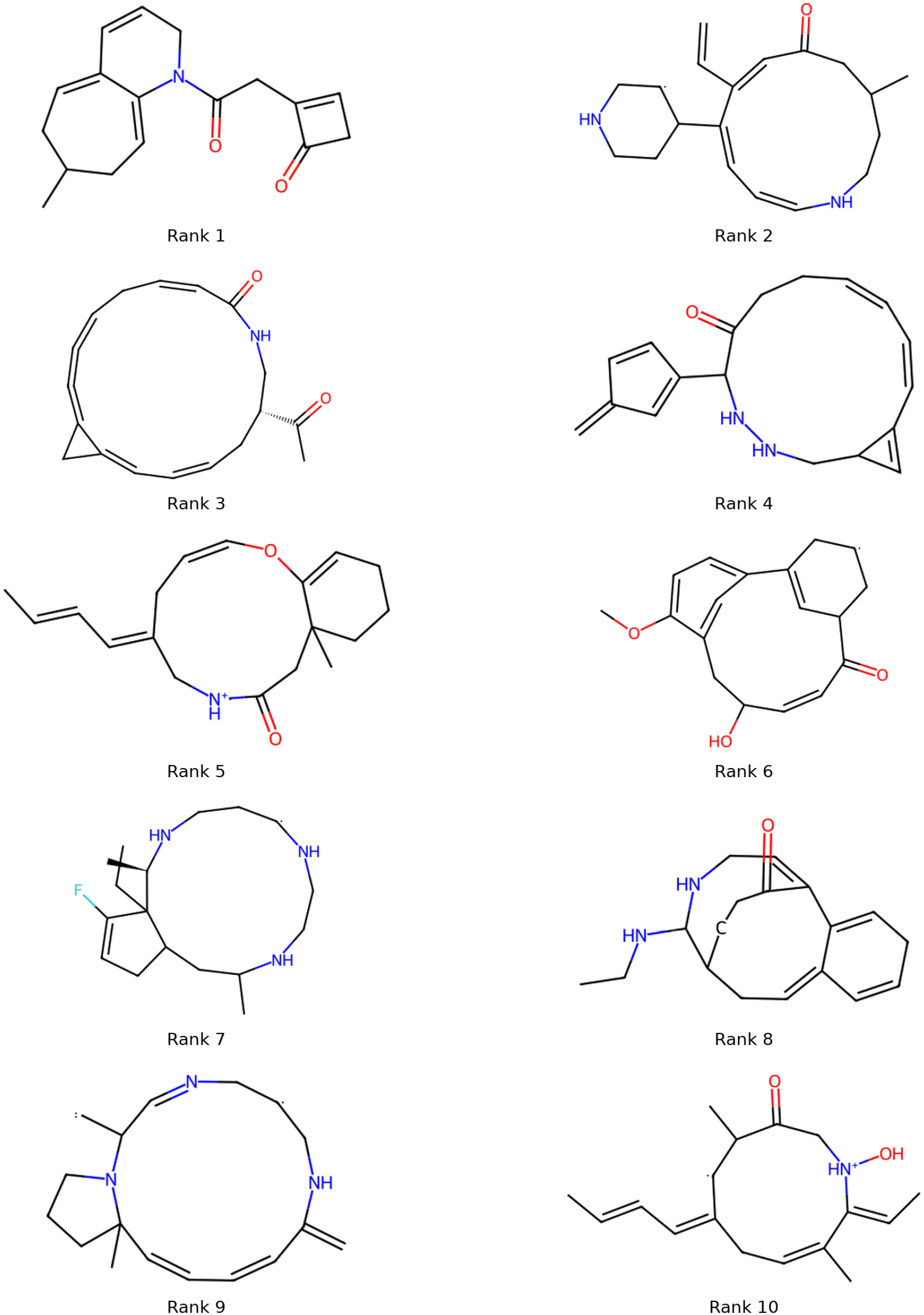
Top 10 DrugDiff candidate molecular structures from Round 1. Candidates are shown in the same rank order as the corresponding top-picks CSV file.

**Figure 6:**
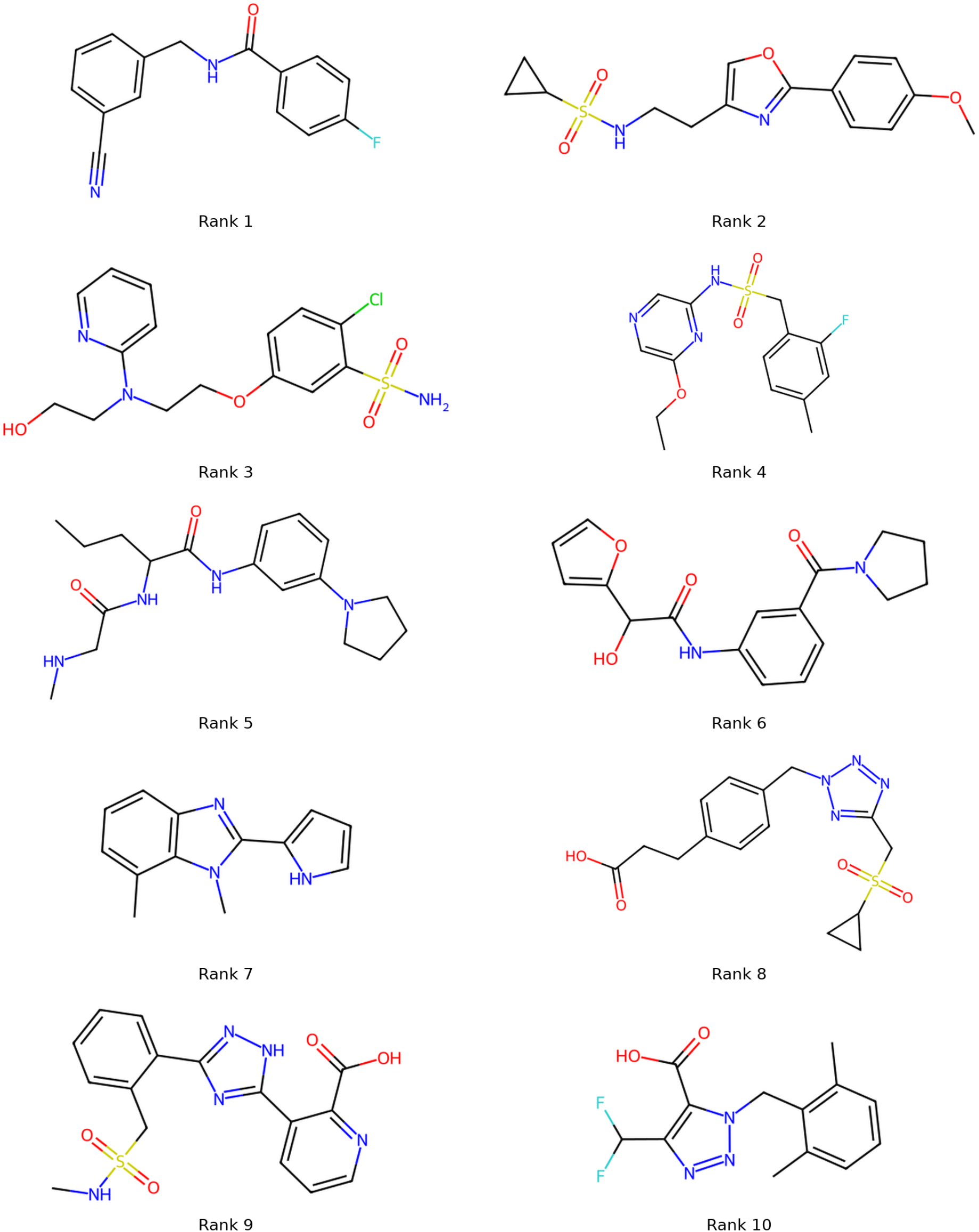
Top 10 BoltzMol-1 candidate molecular structures from Round 2. Candidates are shown in the same rank order as the corresponding top-picks CSV file.

**Figure 7:**
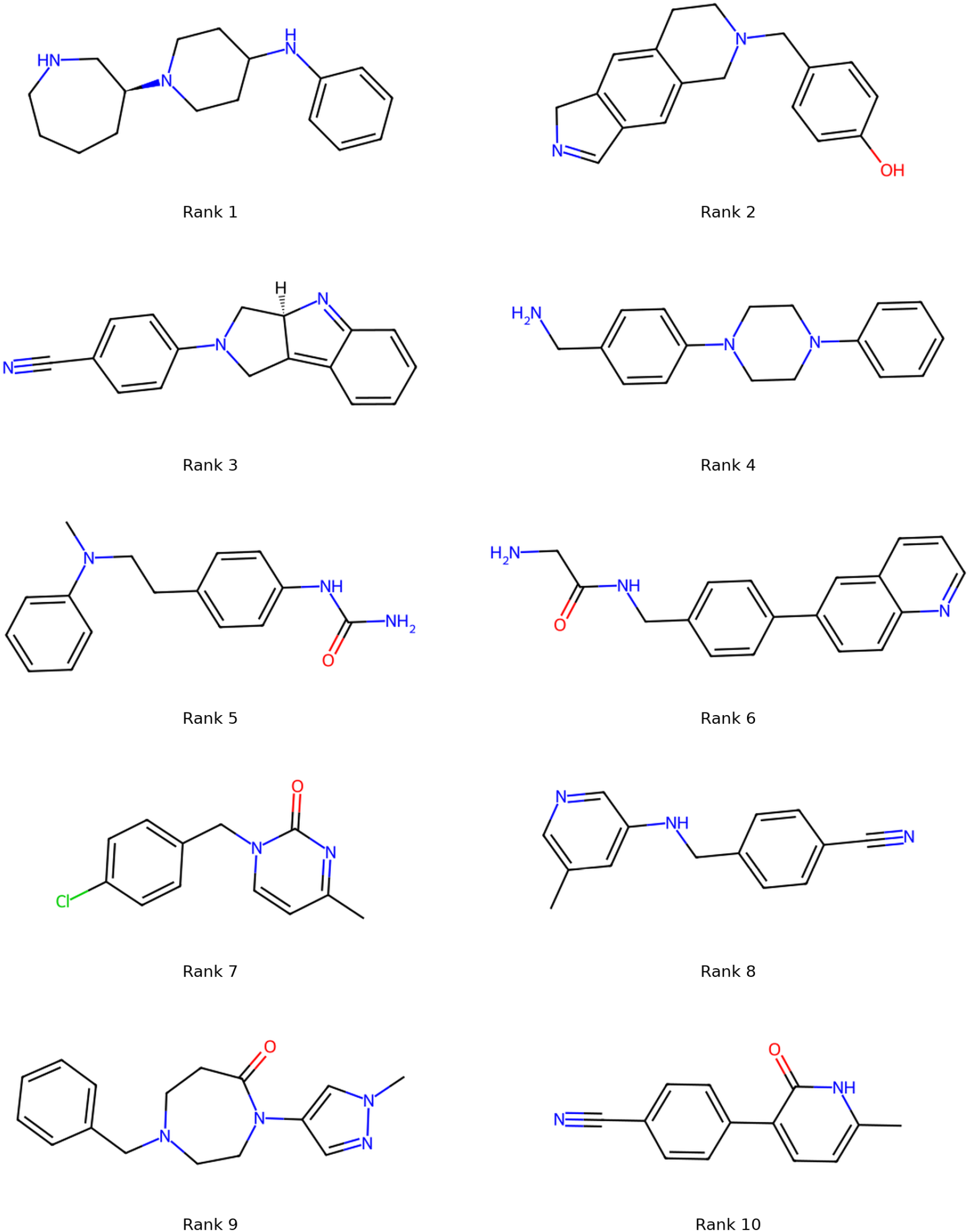
Top 10 FlexFlow candidate molecular structures from Round 2. Candidates are shown in the same rank order as the corresponding top-picks CSV file.

**Figure 8:**
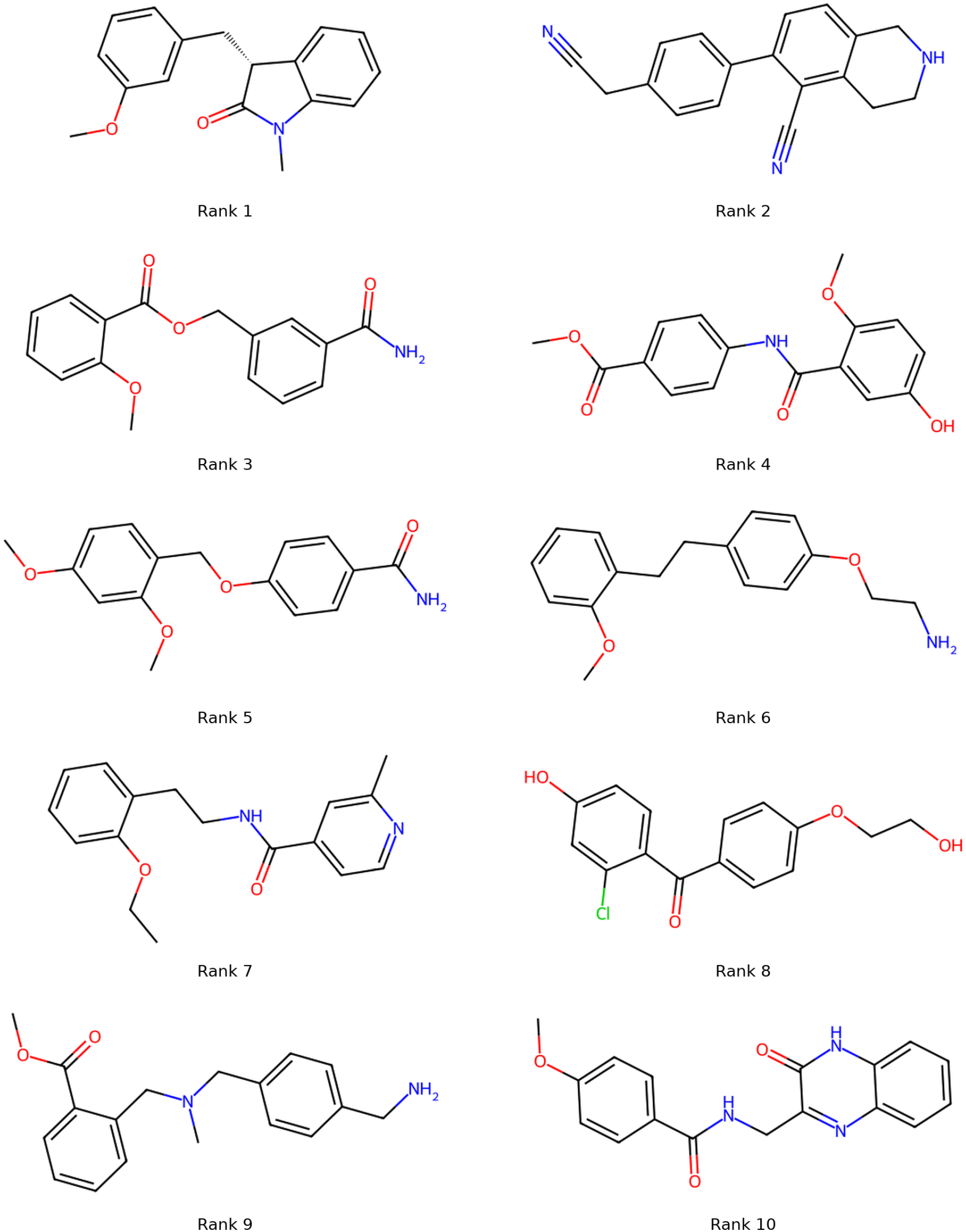
Top 10 DrugFlow-OOD candidate molecular structures from Round 2. Candidates are shown in the same rank order as the corresponding top-picks CSV file.

**Figure 9:**
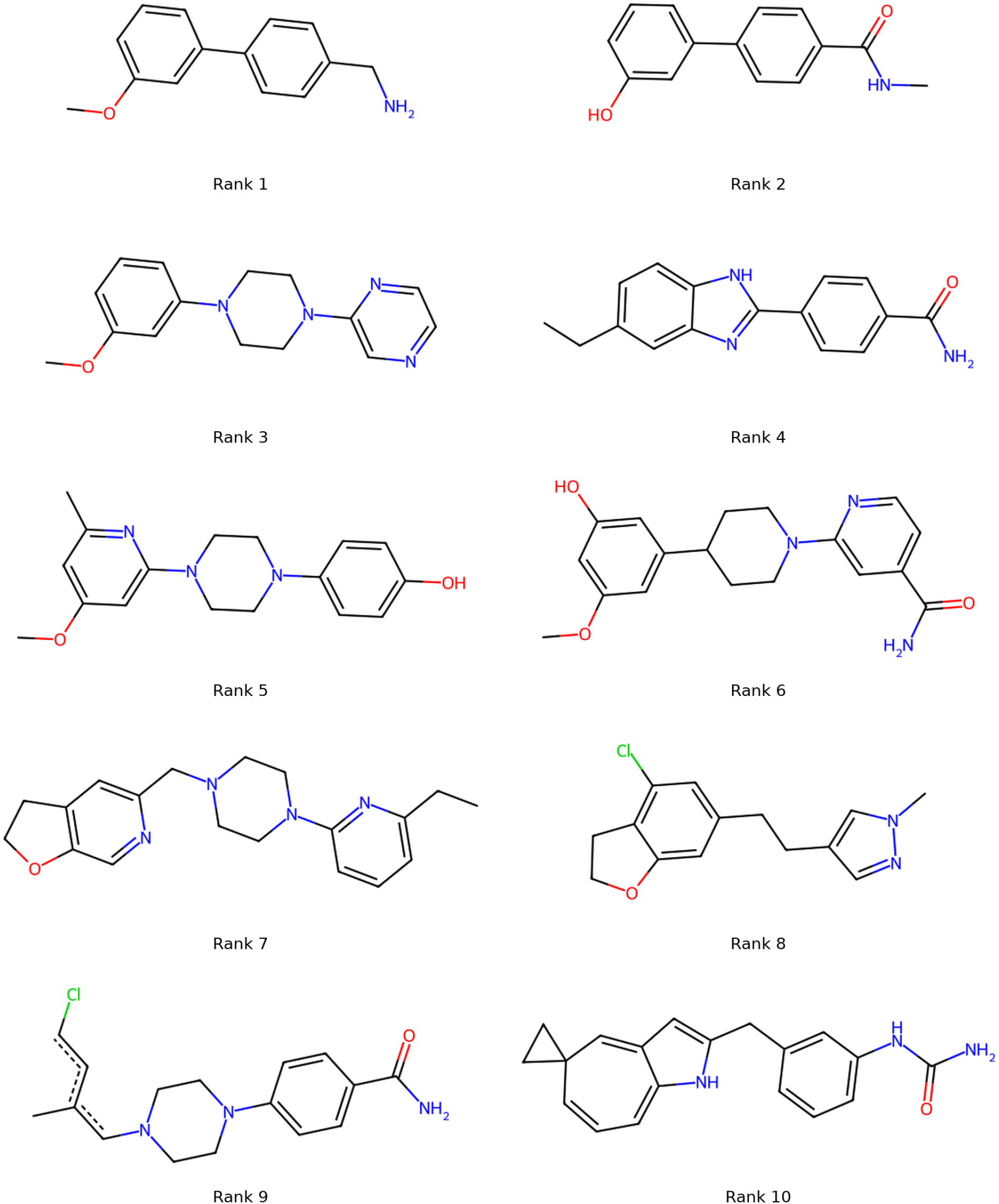
Top 10 DrugFlow-PA candidate molecular structures from Round 2. Candidates are shown in the same rank order as the corresponding top-picks CSV file.

